# Comparing divergence landscapes from reduced-representation and whole-genome re-sequencing in the yellow-rumped warbler (*Setophaga coronata*) species complex

**DOI:** 10.1101/2021.03.23.436663

**Authors:** Stephanie J. Szarmach, Alan Brelsford, Christopher Witt, David P. L. Toews

**Affiliations:** Department of Biology, Pennsylvania State University, State College, PA; Evolution, Ecology, and Organismal Biology Department, University of California Riverside, Riverside, CA; Museum of Southwestern Biology and Department of Biology, University of New Mexico, Albuquerque, NM

**Keywords:** Whole-genome sequencing, reduced-representation sequencing, *F*_ST_, divergence landscapes, mitochondrial introgression

## Abstract

Researchers seeking to generate genomic data for non-model organisms are faced with a number of trade-offs when deciding which method to use. The selection of reduced representation approaches versus whole genome re-sequencing will ultimately affect the marker density, sequencing depth, and the number of individuals that can multiplexed. These factors can affect researchers’ ability to accurately characterize certain genomic features, such as landscapes of divergence—how *F*_ST_ varies across the genomes. To provide insight into the effect of sequencing method on the estimation of divergence landscapes, we applied an identical bioinformatic pipeline to three generations of sequencing data (GBS, ddRAD, and WGS) produced for the same system, the yellow-rumped warbler species complex. We compare divergence landscapes generated using each method for the myrtle warbler (*Setophaga coronata coronata*) and the Audubon’s warbler (*S. c. auduboni*), and for Audubon’s warblers with deeply divergent mtDNA resulting from mitochondrial introgression. We found that most high-*F*_ST_ peaks were not detected in the ddRAD dataset, and that while both GBS and WGS were able to identify the presence of large peaks, WGS was superior at a finer scale. Comparing Audubon’s warblers with divergent mitochondrial haplotypes, only WGS allowed us to identify small (10-20kb) regions of elevated differentiation, one of which contained the nuclear-encoded mitochondrial gene *NDUFAF3*. We calculated the cost per base pair for each method and found it was comparable between GBS and WGS, but significantly higher for ddRAD. These comparisons highlight the advantages of WGS over reduced representation methods when characterizing landscapes of divergence.

## INTRODUCTION

All genomic methods require trade-offs between marker density, sequencing depth, and the degree of multiplexing of individuals (Davey et al. 2011; Toews et al. 2016a). Because high-throughput sequencers generate a fixed amount of data per run and budgets are finite, emphasizing one aspect of these three components usually comes at a cost of the others. Yet, with a large amount of data now generated for a range of biological systems and evolutionary questions, it is now possible to make more specific recommendations for researchers hoping to apply high-throughput analyses to non-model systems.

With high-throughput data, one of the fundamental patterns that researchers aim to quantify between closely related species is how genetic differentiation—particularly *F*_ST_—varies across different regions of the genome (*i*.*e. ‘*genome scans’). These *F*_ST_ ‘divergence landscapes’ were first measured in vertebrates with their application to marine and freshwater three-spined sticklebacks (*Gasterosteus aculeatus*; Hohenlohe et al. 2010), as well as pied flycatchers (*Ficedula hypoleuca*) and collared flycatchers (*Ficedula albicollis*; Ellegren et al. 2012). There is much debate over the interpretation of the peaks in these divergence landscapes, particularly how and whether they relate to an underlying basis for reproductive isolation (i.e. ‘speciation genes’). However, they have proved fruitful for identifying genes underlying important phenotypic differences, such as bill size in Darwin’s finches (Lamichhaney et al. 2015), lateral plates in stickleback (Jones et al. 2012), and pigmentation genes in birds (Toews et al. 2016b; Campagna et al. 2017).

A key consideration for researchers approaching genomic data in non-model systems is: what is the ideal sampling design for estimating useful divergence landscapes? In particular, is it most beneficial to use data from reduced-representation sequencing approaches (i.e. genotype-by-sequencing, GBS, or double-digest restriction site-associated DNA sequencing, ddRAD) or whole-genome re-sequencing (WGS)? Answering this question is challenging, because different methods and approaches are also applied across different biological systems, making generalizations difficult. It would be useful, however, to compare how observed divergence landscapes differ across technologies applied to the same study system. Here we do this among closely related wood warblers.

The yellow-rumped warbler (*Setophaga coronata*) has a long history of molecular study. This species complex is composed of four currently recognized subspecies: *Setophaga coronata coronata*, the myrtle warbler, which breeds in the boreal forest east of the Rocky Mountains and winters in eastern North America, Central America, and the Caribbean; *S. c. auduboni*, the Audubon’s warbler, which breeds west of the Rocky mountains and winters in the southwestern United States, Mexico, and central America; *S. c. nigrifrons*, the black-fronted warbler, which occurs year-round in Mexico; and finally, *S. c. goldmani*, or Goldman’s warbler, which consists of a small population of resident birds confined to Guatemala (Hubbard 1970).

Molecular characterization in *Setophaga coronata* began with Barrowclough (1980), whose analysis of allozyme variation found very few genetic differences between the two main North American subspecies, *S. c. coronata* and *S. c. auduboni*. Brelsford et al. (2009) and Brelsford et al. (2011), using nuclear introns and AFLPs, also supported a high level of genetic similarity, but found the first signs of highly differentiated genomic regions across the group. Brelsford et al. (2011), Milá et a. (2011), and Toews et al. (2014) all later found evidence of mitochondrial introgression—from *S. c. coronata* into *S. c. auduboni—*calling into question previous interpretations of exceptional genetic similarity between these two subspecies based only on mtDNA (Milá et al. 2007). Finally, Toews et al. (2016c) revisited nuclear differentiation with high-throughput reduced-representation sequencing (GBS). This analysis confirmed the genetic diagnosability of all four subspecies, and also identified the first contours of multiple, large (i.e. >1 Mb) highly differentiated genomic regions.

Here, we first compare the heterogeneous divergence landscapes in the yellow-rumped warbler derived from two reduced-representation sequencing methods (GBS and ddRAD), with those obtained from whole genome re-sequencing data (WGS), using an identical bioinformatic pipeline. We compare how these methods differ in their ability to quantify divergence landscapes between the two main forms, *S. c. auduboni* and *S. c. coronata*. Second, we revisit a previous comparison within *S. c. auduboni—* between individuals that are phenotypically indistinguishable but have deeply divergent mtDNA (introgressed-northern versus ancestral-southern clades)—where it was not known if, in addition to mtDNA introgression, portions of the nuclear genome introgressed in conjunction. Past molecular assays using reduced-representation sequencing left open the question of whether there are, in fact, introgressed regions of the nuclear genome that could be identified from higher resolution data. If such nuclear introgression had occurred, we would expect to see regions of differentiation between *S. c. auduboni* individuals that differ in their mitochondrial type.

## MATERIALS AND METHODS

### Comparison datasets

We draw on data from three previously published comparisons of yellow-rumped warblers. The first, from Toews et al. (2016c), used genotype-by-sequencing reads, aligned to the Zebra Finch genome with a coverage averaged across the genome of 0.2X, to assay variation across 37,518 SNPs (for the distributions of coverage across the genome for each dataset see Figs. S1-S3). Briefly, paired-end GBS data was generated by enzymatic complexity reduction with a single restriction enzyme, *PstI*. Second, Toews et al. (2018) used single-end sequencing reads generated from a ddRAD method to assay variation across 19,709 polymorphic SNPs aligned to the first generation of the *Setophaga coronata coronata* genome with an average genome-wide coverage of 0.04X. This method used two restriction enzymes, *SbfI* and *MspI*, to reduce the complexity of the genome. Finally, Baiz et al. (2021) used paired-end sequencing of whole-genomes from randomly sheared, 350 bp size-selected fragments, sequenced to an average coverage of 4X. For the present study, reads from all three datasets were aligned to the new chromosome-level genome assembly of *S. c. coronata* published in Baiz et al. (2021).

To compare the three methods in terms of the cost of raw output, we calculated the cost per base pair for each approach. First, we calculated the number of reads per sequencing lane by multiplying the average reads per individual for each sample included in the current study (a small, but representative, sample of the overall individuals sequenced for the GBS and ddRAD datasets) by the number of individuals multiplexed on a single sequencing lane. We then determined the number of base pairs per lane by multiplying the number of reads by the run length (100 or 150bp) and by a factor representing the sequencing chemistry (x1 for single-end or x2 for paired-end). Lastly, we divided the cost per lane at the time of sequencing by this estimated total output per lane (recognizing that costs will be variable across institutional sequencing centers, but generally comparable), resulting in an estimate of dollars per base pair.

### Sampling information

We assessed genome-wide *F*_*ST*_ between *S. c. coronata* and two groups of *S. c. auduboni* from divergent mitochondrial lineages. *S. c. auduboni* in the southernmost portion of their range (Arizona and New Mexico) possess mitochondrial haplotypes belonging to the same clade as those of *S. c. nigrifrons* (hereafter referred to as “southern haplotypes”; Brelsford et al. 2011, Toews et al. 2014). In contrast, *S. c. auduboni* north of this region share mtDNA haplotypes with *S. c. coronata* (hereafter “northern haplotypes”), likely resulting from past mitochondrial introgression between these taxa. In addition to examining the landscape of divergence between *S. c. coronata* and *S. c. auduboni*, we also survey patterns of *F*_*ST*_ between *S. c. auduboni* individuals with northern mitochondrial haplotypes and those with southern haplotypes. For the WGS and GBS analyses, we used sequences obtained from five *S. c. coronata* individuals, five *S. c. auduboni* individuals with southern mitochondrial haplotypes, and five *S. c. auduboni* individuals with northern haplotypes (Table 1). While the same sampling scheme was used for both analyses, each dataset contains different individuals sequenced as part of previous studies (GBS samples from Toews et al. 2016c; WGS samples from Baiz et al. 2021; Figure 1). The ddRAD analysis used five *S. c. coronata* samples and five *S. c. auduboni* with northern mtDNA haplotypes (Toews et al. 2018).

**Table 1.** Subspecies identification, mitochondrial clade (N – northern, S – southern), and collection location of samples from three datasets generated using whole-genome re-sequencing (wgs), genotype-by-sequencing (gbs), or double-digest restriction site-associated DNA sequencing (ddrad).

| Datatype | Sample ID | Subspecies | mtDNA Haplotype | Latitude | Longitude |
| --- | --- | --- | --- | --- | --- |
| wgs | CUMV4915 | coronata | N | 44.25136 | -73.959019 |
| wgs | QF11T02 | coronata | N | 43.802827 | -74.651096 |
| wgs | QF10T03 | coronata | N | 43.718615 | -74.784205 |
| wgs | QF09T03 | coronata | N | 43.689824 | -74.751738 |
| wgs | QF11T06 | coronata | N | 43.771995 | -74.637935 |
| wgs | MSB41238 | auduboni | N | 36.0958 | -108.8874667 |
| wgs | MSB41230 | auduboni | N | 36.24998333 | -109.0467 |
| wgs | MSB40772 | auduboni | N | 36.10418333 | -108.88745 |
| wgs | MSB40787 | auduboni | N | 36.24998333 | -109.0467 |
| wgs | MSB44117 | auduboni | N | 36.08096667 | -108.8656167 |
| wgs | MSB40770 | auduboni | S | 36.09616667 | -108.8883333 |
| wgs | MSB41418 | auduboni | S | 36.89825 | -108.88745 |
| wgs | MSB41233 | auduboni | S | 36.07993333 | -108.8837167 |
| wgs | MSB40788 | auduboni | S | 36.07993333 | -108.8837167 |
| wgs | MSB40775 | auduboni | S | 36.07961667 | -108.8829167 |
| gbs | 03n1310 | coronata | N | 45.161725 | -69.343091 |
| gbs | 03n1312 | coronata | N | 45.161725 | -69.343091 |
| gbs | 03n1313 | coronata | N | 45.161725 | -69.343091 |
| gbs | 03n1319 | coronata | N | 45.161725 | -69.343091 |
| gbs | 03n1321 | coronata | N | 45.161725 | -69.343091 |
| gbs | ED24A05 | auduboni | N | 52.4929 | -121.71198 |
| gbs | ED25A02 | auduboni | N | 52.46222 | -121.68444 |
| gbs | ED25A03 | auduboni | N | 52.49645 | -121.71039 |
| gbs | ED26A03 | auduboni | N | 52.5055 | -121.698 |
| gbs | ED26A05 | auduboni | N | 52.50343 | -121.69787 |
| gbs | JE14T01 | auduboni | S | 34.986719 | -111.46873 |
| gbs | JE28T03 | auduboni | S | 36.641838 | -112.16567 |
| gbs | KF02T04 | auduboni | S | 38.610871 | -111.66817 |
| gbs | JF16T05 | auduboni | S | 35.889229 | -106.63446 |
| gbs | JF22T03 | auduboni | S | 37.986651 | -106.54618 |
| ddrad | EE22A03 | coronata | N | 54.58789 | -110.27156 |
| ddrad | EE25A04 | coronata | N | 54.72454 | -110.0651 |
| ddrad | GF19A09 | coronata | N | 61.15445 | -149.73282 |
| ddrad | GF20A06 | coronata | N | 61.15942 | -149.74254 |
| ddrad | GF21A10 | coronata | N | 61.15523 | -149.75172 |
| ddrad | GE26A02 | auduboni | N | 49.16708 | -121.33685 |
| ddrad | GE26A04 | auduboni | N | 49.15248 | -121.31281 |
| ddrad | GE26A05 | auduboni | N | 49.20206 | -121.36902 |
| ddrad | GE26A06 | auduboni | N | 49.20591 | -121.36852 |
| ddrad | GE26A08 | auduboni | N | 49.22609 | -121.3794 |

**Figure 1.**
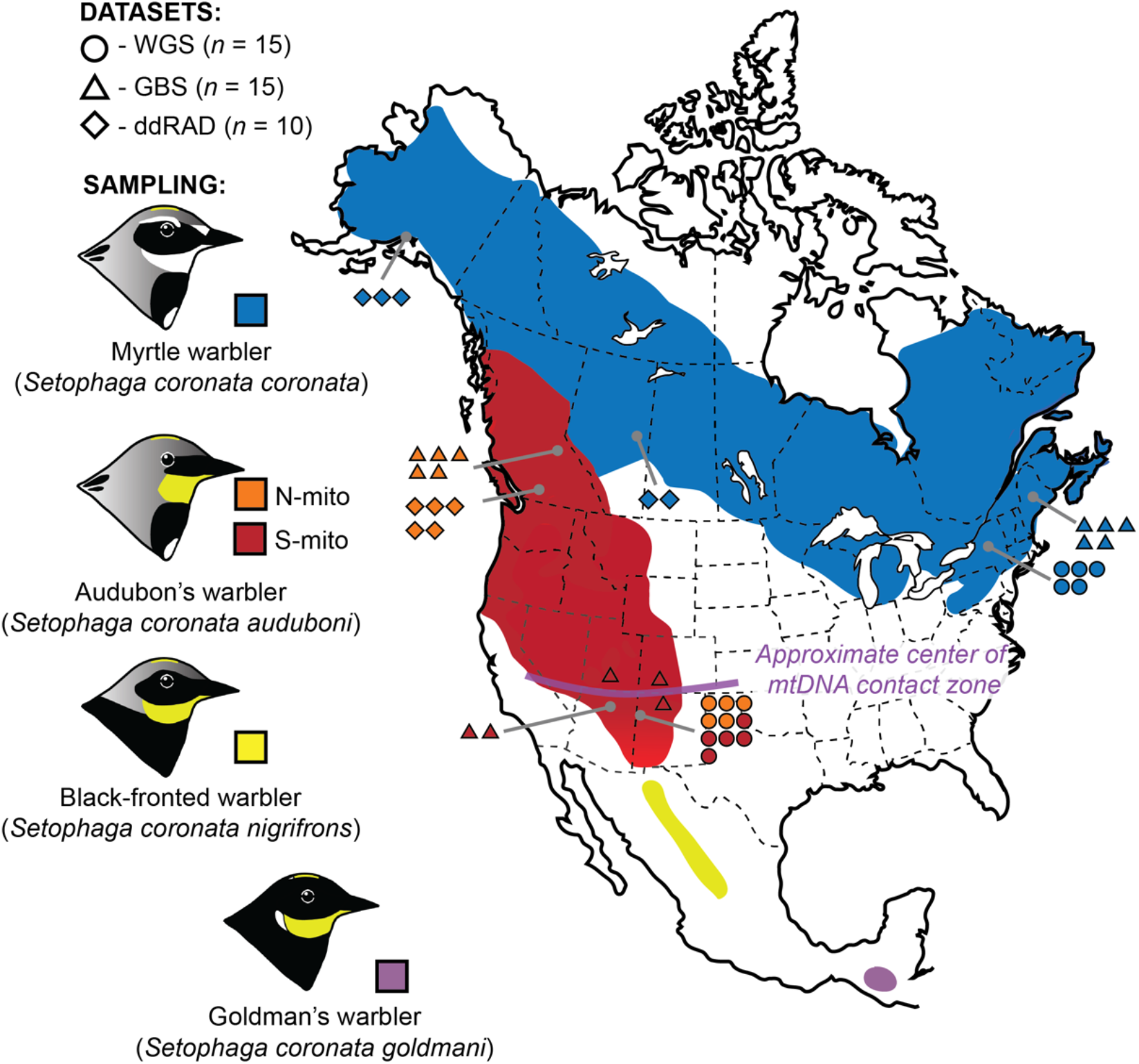
Sampling map illustrating all four subspecies in the yellow-rumped warbler species complex, including sampling locations for the samples used in the present study. *S. c. nigrifrons* and *S. c. goldmani* are shown for reference, but were not included in the present study.

### Bioinformatic analyses

We first used the program AdapterRemoval (Schubert et al. 2016) to trim and collapse overlapping paired reads with the following options: “-collapse -trimns -minlength 20 -qualitybase 33”. To align reads from each dataset to the *S. c. coronata* reference genome, we used BowTie2 (Langmead and Salzberg 2012) with the “very-sensitive-local” presets and the “-X” option, which assigns the maximum fragment length allowed for valid paired-end alignments, set to 700 bp for the WGS and GBS data (ddRAD reads were single-end). PCR duplicates were marked with PicardTools version 2.20.8 (https://broadinstitute.github.io/picard/); we only excluded duplicates from WGS analyses (Andrews et al. 2016). We assessed sequence quality and coverage for the alignments using Qualimap version 2.2.1 (Okonechnikov 2016).

For the WGS and GBS datasets, we estimated *F*_*ST*_ across the genome between three population pairs: *S. c. coronata* vs. *S. c. auduboni* – northern haplotype; *S. c. coronata* vs. *S. c. auduboni* – southern haplotype; and *S. c. auduboni* – northern haplotype vs. *S. c. auduboni* – southern haplotype. As the ddRAD dataset did not include individuals with the southern haplotype, for this analysis we only estimated *F*_*ST*_ between *S. c. coronata* and *S. c. auduboni* – northern haplotype. For all datasets, *F*_*ST*_ was estimated using ANGSD version 0.929 (Korneliussen et al. 2014), which utilizes genotype likelihoods to account for uncertainty in low-coverage sequence data. First, we calculated the sample allele frequencies for each population using the -doSaf command. Using these allele frequency likelihoods, we generated two-population site frequency spectrums and estimated *F*_*ST*_ for each population pair using the realSFS program. We calculated *F*_*ST*_ across non-overlapping 10-kb windows using the fst stats2 option in realSFS with the options -win and -step both set to 10,000. The results of the sliding window analysis were visualized using the R package qqman (Turner 2018).

To contrast the three methods in terms of information content, we used the output of the sliding window analysis to estimate the number of “data sites” in each. As ANGSD does not use called genotypes to estimate allele frequencies—instead using genotype likelihoods—we cannot make straightforward comparisons of the raw number of SNPs produced with each method. Instead, the pipeline outputs the number of informative sites where there is associated data, whether or not the site is polymorphic across samples or has high missingness. To compare the patterns of *F*_ST_ across the three methods, we matched common 10 kb and 1 mb windows, and used a Pearson correlation in R 3.5.1 (R Core Team 2018). We also used this estimate of informative data sites as another comparison of the dollar value among the methods. We multiplied the cost per individual by ten, as all the *F*_ST_ included five individuals from two populations. We then divided this cost by the number of data sites, and transformed this to cost per mb.

We used principal component analysis to visualize genomic variation in the differentiated region on chromosome 12 (only WGS data; see below). To do this, we used the genotype likelihoods to estimate covariance matrices with PCAngsd 0.981 (Meisner and Albrechtsen 2018) using default settings. We then used the eigen function in R to compute and plot the first 2 eigenvectors.

## RESULTS

The average number of reads varied significantly across the three datasets, with average reads totaling 0.6 million per individual for ddRAD, 4.8 million per individual for GBS, and 31.0 million per individual for WGS. Across the three different library preparation methods, WGS and GBS have comparable value when comparing costs per bp of sequence generated, with both performing better than the ddRAD approach (Table 2). We found that both GBS and WGS produced 2.5X more sequence per dollar than ddRAD, even though the ddRAD project was highly multiplexed. In terms of information content, the three approaches produced predictable numbers of informative sites per 10 kb window: ddRAD, average 141 data sites / 10 kb; GBS, average 364 data sites / 10 kb, and WGS, 9361 data sites / 10 kb.

**Table 2.** Results of the cost comparison between genotype-by-sequencing (GBS), double-digest restriction site-associated DNA sequencing (ddRAD), and whole-genome re-sequencing (WGS). Presented for each method are the number of individuals multiplexed on a sequencing lane, the sequencing technology and chemistry, length in base pairs of the sequencing run, the average reads produced per individual, cost per lane, individual, and megabase (in U.S. dollars), the number of informative data sites used in the *F*_*ST*_ analysis, and the cost (USD) per megabase of informative sites for an analysis that included *n* = 10 individuals.

| Library method | Multiplex | Technology | Sequencing Year | Chemistry | Run length | Average reads per individual | Cost per lane (USD) | Cost per individual | Cost per megabase | Informative data sites for $F_{ST}$ | Cost per data site (MB) |
| --- | --- | --- | --- | --- | --- | --- | --- | --- | --- | --- | --- |
| GBS | 96 | HiSEQ 2000 | 2014 | paired-end | 100 | 4,796,893 | 2,362 | 25 | 0.051 | 26,027,347 | 9.5 |
| ddRAD | 192 | HiSEQ 2500 | 2016 | single-end | 100 | 607,597 | 1,350 | 7 | 0.127 | 2,825,361 | 24.9 |
| WGS | 24 | NextSeq 500 | 2018 | paired-end | 150 | 31,034,023 | 4,791 | 200 | 0.043 | 948,976,685 | 2.1 |

Only GBS and WGS were able to capture the heterogeneous landscape of divergence between myrtle and Audubon’s warblers (Figure 2), whereas ddRAD was not able to identify clear *F*_*ST*_ peaks. Comparing GBS and WGS, while GBS data could clearly identify the presence or absence of large (i.e. 1 Mb) *F*_*ST*_ peaks, only WGS data adequately identified the clear contours of the start and endpoints of these peaks (Figure 3), as well as their graded shoulders, and subtle reductions within a peak (Figure 3c).

**Figure 2.**
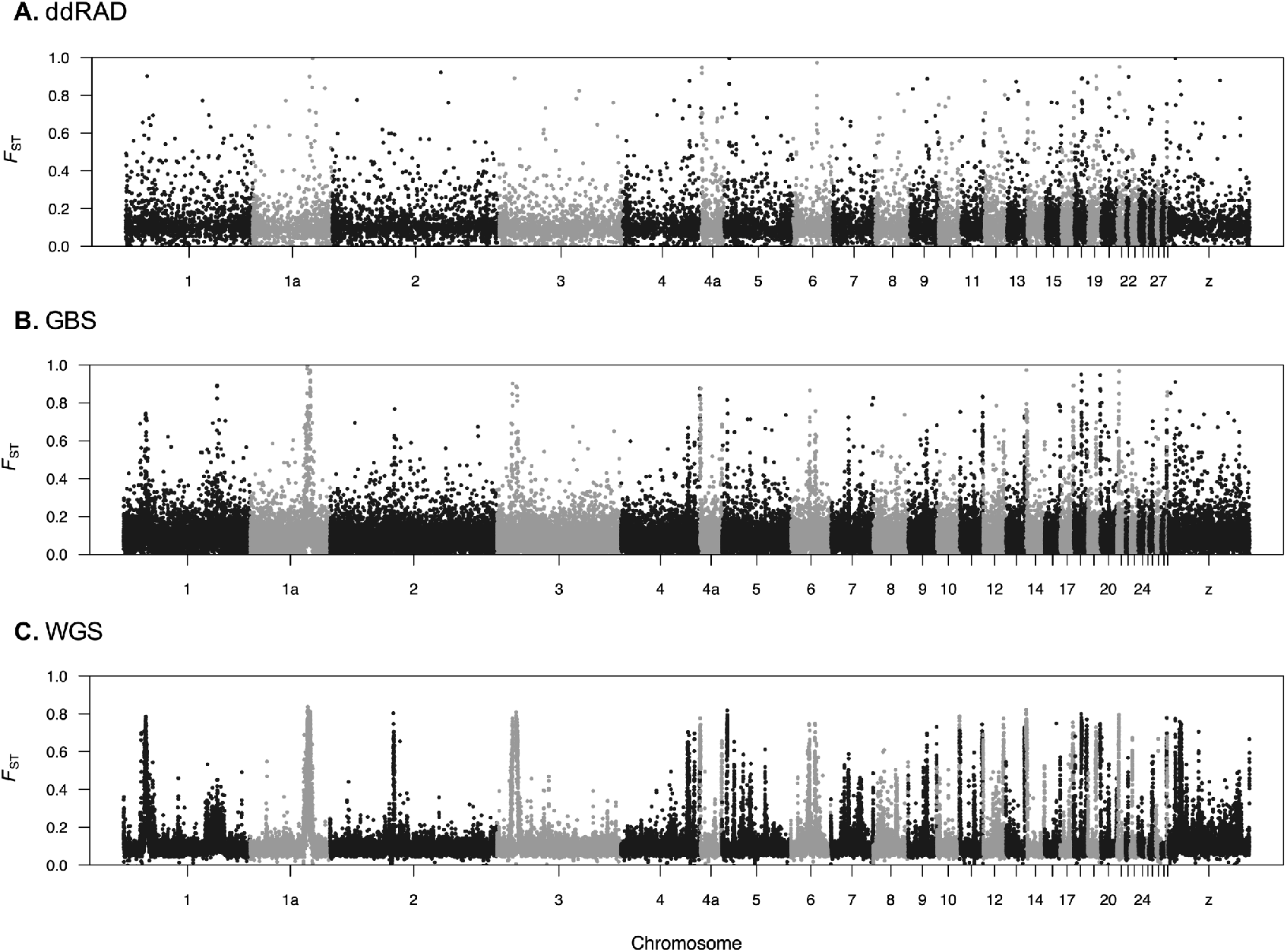
Genome-wide *F*_*ST*_ for *S. c. coronata* vs. *S. c. auduboni* estimated in 10 kb windows across the genome using data from (A) double-digest restriction-site associated DNA sequencing, (B) genotyping by sequencing, and (C) whole-genome re-sequencing.

**Figure 3.**
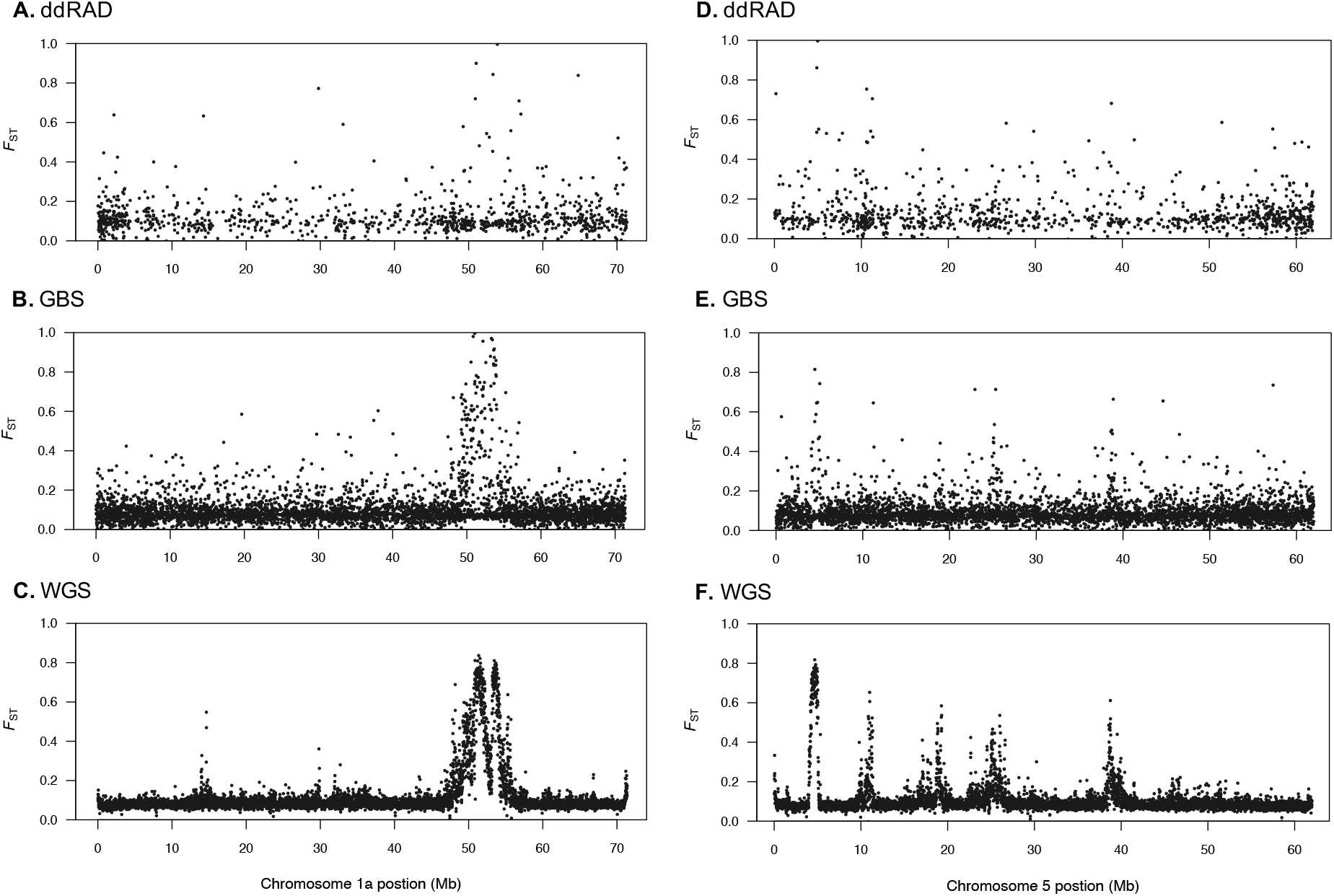
Detail of *F*_*ST*_ estimated between *S. c. coronata* and *S. c. auduboni* in 10 kb windows over chromosome 1a (A-C) and chromosome 5 (D-F) using double-digest restriction-site associated DNA sequencing (A,D), genotyping by sequencing (B,E), and whole-genome re-sequencing (C,F) datasets.

Correlations between common windows were overall low across the three methods (Figure 4). This was particularly true for comparisons of common 10 kb windows (Figure 4a-c; Pearson’s correlation: WGS / GBS = 0.32, GBS / ddRAD = 0.11, WGS / ddRAD = 0.11), where WGS showed that in many of the windows that were found to have low *F*_*ST*_ in the GBS and ddRAD datasets *F*_ST_ is likely much higher (i.e. the “L-shaped” distribution of points). At a larger scale (1 mb windows; Figure 4 d-f) the correlations across methods were higher (Pearson’s correlation: WGS / GBS = 0.78, GBS / ddRAD = 0.39, WGS / ddRAD = 0.41), with the comparison between GBS and WGS showing the highest concordance at this scale.

**Figure 4.**
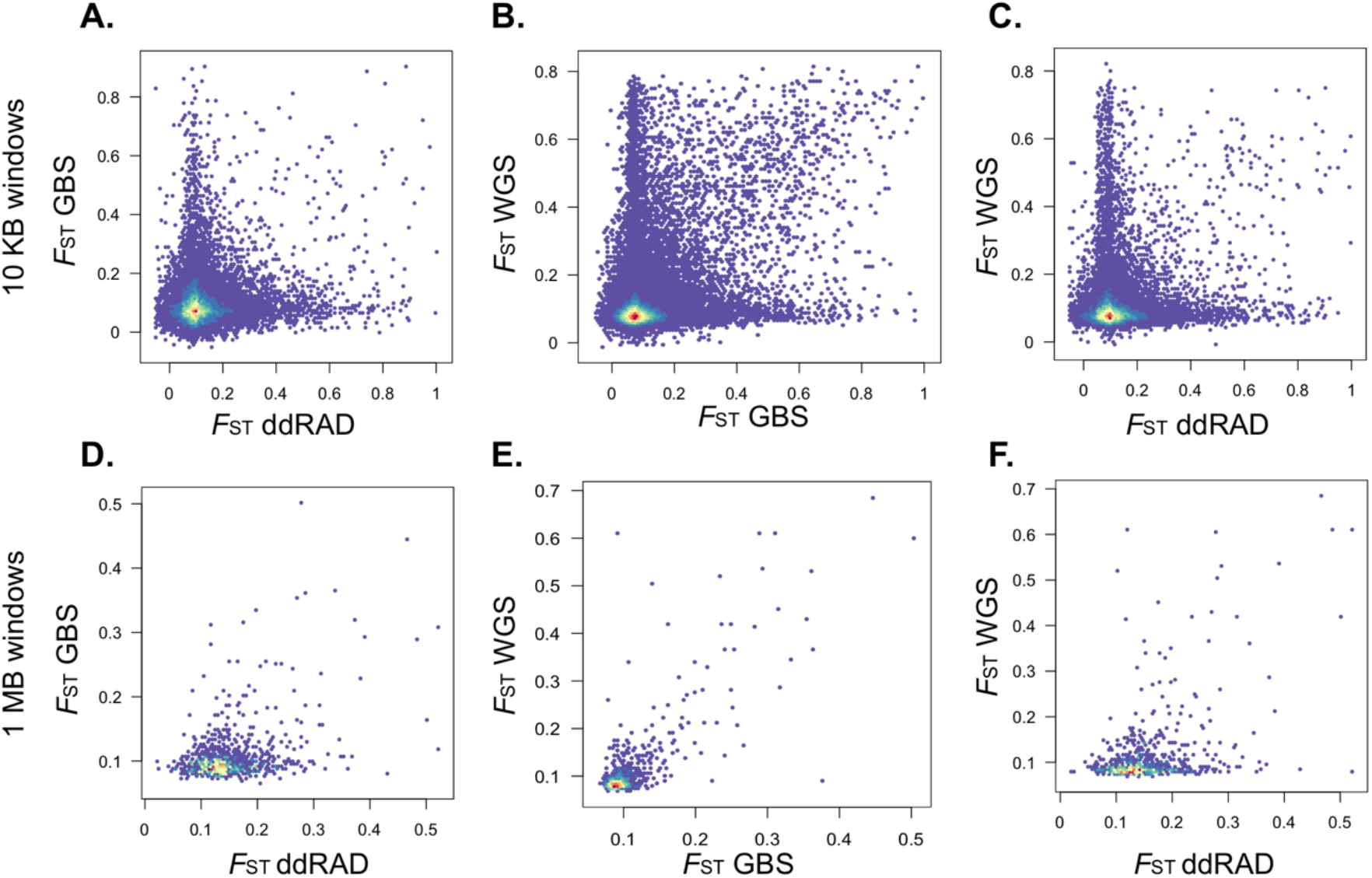
Correlations between common windows of the genome-wide *F*_ST_ analysis (for *S. c. coronata* vs. *S. c. auduboni*) across different sequencing methods: double-digest restriction-site associated DNA sequencing (A, D), genotyping by sequencing (B, E), and whole-genome re-sequencing (C, F). Correlations were calculated for 10 kb windows (A-C) and 1 MB windows (D-F).

Comparing Audubon’s warblers from the two different mitochondrial clades (northern versus southern haplotypes), both GBS and WGS exhibited very low genome-wide *F*_*ST*_, as found in prior studies. The average genome-wide *F*_*ST*_ was 0.024 for GBS and 0.026 for WGS data. However, the WGS data showed five windows with slightly elevated divergence (*F*_*ST*_ > 0.2) that had not been previously identified using reduced representation sequencing, occurring on chromosomes 2 (one 10 kb window), 9 (two 10 kb windows), and 12 (two 10 kb windows; Figure 5A). Notably, the *F*_*ST*_ windows on chromosomes 9 and 12 are adjacent windows in the genome, an observation very unlikely due to chance alone. While GBS data had several moderate *F*_ST_ windows, none of the most differentiated windows (top 0.1%) occur in adjacent windows.

**Figure 5.**
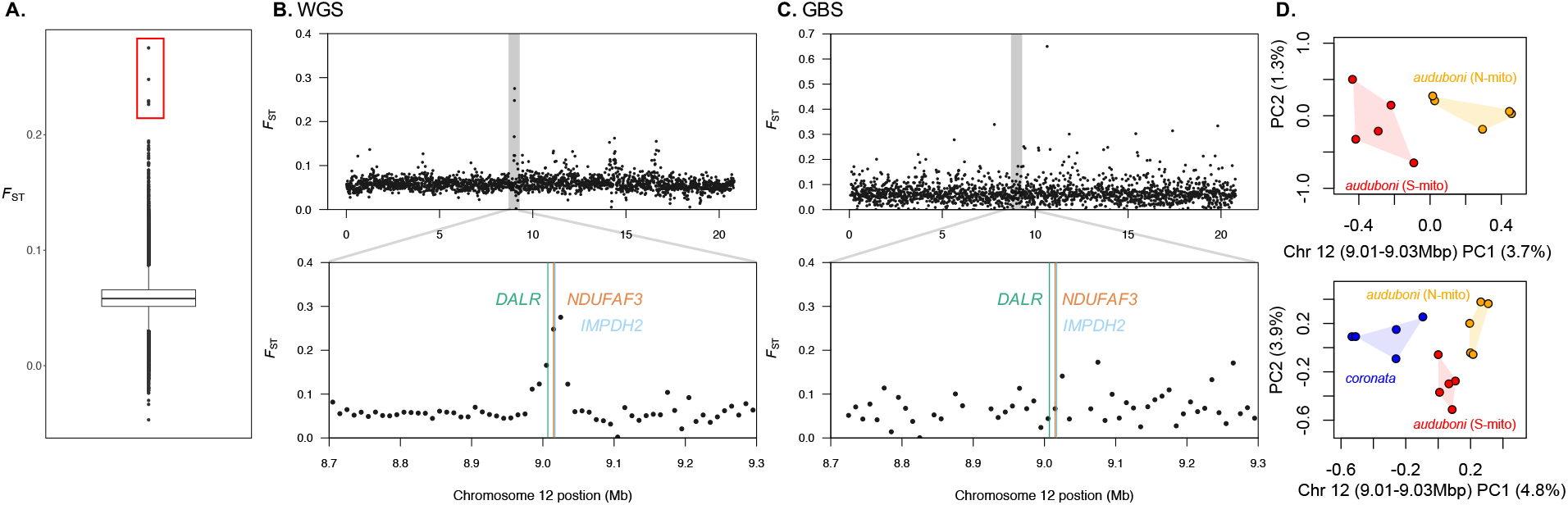
(A) Distribution of *F*_*ST*_ values for 10 kb windows between *S. c. auduboni* from the northern mtDNA clade vs. *S. c. auduboni* from the southern mtDNA clade. The red box highlights the windows with elevated divergence that were examined further. *F*_*ST*_ across chromosome 12 calculated in 10 kb windows for *S. c. auduboni* from the northern mtDNA clade vs. *S. c. auduboni* from the southern mtDNA clade using (B) whole genome sequencing and (C) genotyping by sequencing. The grey box highlights the peak of elevated divergence identified in the WGS data, and the panel beneath exhibits a detailed view of this region. The colored lines indicate the positions of three genes (DALR, NDUFAF3, and IMPDH2) located within this region of divergence. (D) Principal components analysis of variation within the divergent region on chromosome 12 from whole genome sequencing data. The top plot shows *S. c. auduboni* -northern mtDNA haplotype and *S. c. auduboni* -southern mtDNA haplotype alone, while the bottom plot includes *S. c. coronata*.

Examining the annotation of the *S. c. coronata* reference genome found that three genes are located within the divergent region on chromosome 9: GP5, LRRC15, and CPN2. Three genes were also located in the divergent region on chromosome 12: DALR, IMPDH2, and NDUFAF3 (Figure 5B, C). No genes were found within the divergent region on chromosome 2.

We focused on the region on chromosome 12, as it contains the only nuclear-mitochondrial gene (*NDUFAF3*). From the WGS dataset, in the PCA for this region (chromosome 12, between 9,010,000-9,030,000 bp), PC1 clearly clustered *S. c. auduboni* warblers with different mitochondrial types. PC1 also explained disproportionately more variation than all other PC axes (percent of variance explained PC1: 3.7, PC2: 1.3, PC3: 1.1, PC4: 0.8). In a separate PCA, also including *S. c. coronata* individuals, PC1 separated *S. c. coronata* from all *S. c. auduboni* individuals. PC2 then separated *S. c. auduboni* individuals by mitochondrial clade, with *S. c. coronata* clustering with *S. c. auduboni* with the northern haplotypes along PC2. In this analysis, both PC1 and PC2 explain disproportionately more variation than all other PC axes (percent of variance explained PC1: 4.8, PC2: 3.9, PC3: 1.3, PC4: 1.1).

## DISCUSSION

With three generations of genome sequencing methods applied to the same biological system— different subspecies of the yellow-rumped warblers—using the same bioinformatic pipeline, we have provided an empirical contrast of the different tradeoffs for reduced-representation (ddRAD and GBS) versus whole genome re-sequencing (WGS). We have done this in the context of quantifying landscapes of genomic divergence (*F*_*ST*_), which is of broad interest to studies of molecular ecology.

### Contrasting methods for quantifying divergence landscapes

Consistent with previous studies (Toews et al. 2016c; Irwin et al. 2018; Baiz et al. 2021), contrasts between *S. c. coronata* and *S. c. auduboni* revealed a highly heterogeneous landscape of relative divergence, as measured by *F*_ST_. This landscape consists of fixed—or nearly fixed—SNPs clustered into large 1-2 Mb regions. As has been suggested in other avian systems (Battey et al. 2020), these large divergence peaks are likely the result of reduced recombination and linked selection (Burri et al. 2016). In several cases, these regions could be associated with centromeres (e.g. Ellegren et al. 2012)—chromosomal features with significantly reduced recombination—although we currently have no direct knowledge of their genomic locations in the warbler genome. However, as the handful of distinct divergent regions across chromosome 5 illustrate (Figure 3F), in several cases the divergent regions between these warblers are more widely distributed across the chromosome, in contrast to what would be expected from a single centromere location. Thus, it is likely these peaks are the result of other processes reducing local recombination rates and contributing to elevated divergence.

In terms of information value, we find that WGS, even with a small number of individuals and low-to-moderate coverage, provides substantially higher resolution genome scans compared to both reduced-representation approaches (Figure 2). Correlations between *F*_*ST*_ of common windows were low across sequencing methods when using smaller (10kb) window sizes, suggesting that GBS and ddRAD are not capturing regions of divergence that were detected using WGS. Cross-method correlations of *F*_*ST*_ for larger (1MB) windows were higher, indicating that the methods characterize *F*_*ST*_ more consistently at a coarser scale. The reduced representation approaches were much less effective at identifying regions with elevated *F*_*ST*_ at the fine scale and do not provide the same detailed picture of the landscape of divergence that is found using WGS.

We found that the cost per base pair for WGS was comparable to, if not better than, both GBS and ddRAD (Table 2). Some of this reduction in cost effectiveness for the ddRAD approach may stem from sequencing a smaller proportion of the genome at higher depth. The benefit of this is an increased ability to accurately call genotypes for specific SNPs, but with the tradeoff that a smaller number of informative sites will be sequenced with more redundancy, leading to a higher cost per base pair. While the motivation behind many reduced representation sequencing approaches is overall cost reduction at the expense of genomic coverage, without some background knowledge of differentiation patterns, regions of divergence may be too small or few to be captured by these methods, leading to inconclusive results. For example, large ddRAD datasets were generated to identify *F*_ST_ divergence peaks among *Sporophila* seedeaters (Campagna et al. 2015), *Vermivora* warblers (Toews et al. 2016b), and *Colaptes* woodpeckers (Aguillon et al. 2018). In each case, ddRAD failed to detect any fixed or highly divergent SNPs across taxa. Small peaks were then subsequently recovered by using WGS data (e.g. Campagna et al. 2017; Aguillion et al. 2020), leaving little value for the time and expense invested in generating ddRAD data. Thus, we have found that in practice, we would recommend in situations without prior knowledge of background differentiation, researchers wishing to quantify landscapes of divergence in *F*_ST_ move straight to WGS data instead of first applying reduced representation methods.

Enzymatically reduced DNA sequencing methods do provide flexibility in the proportion of the genome sequenced and the depth of sequencing, mostly depending on the number of restriction enzymes (i.e. single versus double) and their frequency of cutting. The current comparison is limited by the enzymes selected during previous projects, which do not produce the highest marker densities possible using ddRAD or GBS. Adjusting methods to increase marker density may allow ddRAD and GBS to provide a much better approximation of genome-wide patterns of divergence. However, it can be challenging to predict how many markers will be obtained using a given set of restriction enzymes for non-model species without an available reference genome, and this kind of incremental fine-tuning can therefore increase costs. For systems without existing genomic resources, researchers may be better served by an initial investment in low-coverage WGS, saving both in terms of cost per informative data site, and in the time required to optimize a reduced representation method. This is because the main determinant of depth and coverage in WGS is genome size and ploidy, which have both been estimated for a large number of plant and animal species (e.g. Pellicer and Leitch 2020, Gregory 2021).

One clear benefit of using WGS data from a handful of individuals sampled per population is that researchers can take advantage of tool sets from the realms of both population genetics as well as phylogenetics. For example, using a broader dataset of *Setophaga* warblers—including the yellow-rumped warbler samples used in the present study—Baiz et al. (2021) used an identical sampling design as applied here, generating 4-5X WGS for five or more individuals per species. Baiz et al. (2021) was able to collapse the data to extract sequence data for phylogenetic inference—*e*.*g*. to compare pigmentation gene trees to the ultra conserved element species tree—and also utilize population genetic analyses from genotype likelihoods to estimate *F*_ST_, population branch statistics, and introgression statistics such as ABBA-BABA. Thus, this approach of having a single, high quality chromosome level assembly for the family, with low coverage WGS data aligned for multiple individuals of each species, is likely to produce important evolutionary insights at significantly reduced cost, compared to reference-quality assemblies for each species. An important caveat, however, is that the majority of the data in this approach is currently generated using short-read sequencing, which is less able to capture large, structural variants among populations.

It is likely that the difference in per base pair sequence cost between the reduced-representation and WGS approaches is affected more by sequencing platform than by library preparation method. Yet, enzymatic complexity reduction can have unexpected downstream effects on usable data (reviewed in Andrews et al. 2016). For example, in the present study, due to the low complexity of the ddRAD libraries, we required a spike-in of >5% PhiX control, which reduced the number of sample reads for this project. More generally, however, reduced-representation approaches are clearly important for many applications where fine genomic resolution is not required, such as estimating genome-wide hybrid ancestry (Toews et al. 2018, Walsh et al. 2020), paternity (Thrasher et al. 2018), genomic diversity (Nyinondi et al. 2020), and population structure (Lavretsky et al. 2019).

### Whole-genome analysis of mitochondrial introgression

While GBS and WGS produced qualitatively similar results at broad (i.e. 1 Mb) genomic scales, particularly between the nominate forms *S. c. coronata, and S. c. auduboni* (Figure 4), only WGS data allowed us to identify even smaller regions of elevated divergence (approximately 20 kb) within Audubon’s warblers that differ in mitochondrial backgrounds. Previous studies have found significant mito-nuclear discordance within the yellow-rumped warbler species complex that likely resulted from asymmetric introgression of mitochondria from *S. c. coronata* into *S. c. auduboni* (Milá et al. 2011, Brelsford et al. 2011, Toews et al. 2014). This prior work showed that the introgressed mitochondrial types were broadly associated with a shift in migratory behavior, with the gradual increase in migratory propensity in Audubon’s warblers—inferred indirectly by stable isotopes—coincident with the shift in mitochondrial types (Toews et al. 2014). Moreover, mitochondrial genes encoding NADH proteins in complex I had many more fixed amino acid substitutions between Audubon’s warblers with northern vs. southern mtDNA haplotypes than genes encoding proteins in the other complexes, and these changes likely translated into a small but significant difference in mitochondrial respiratory efficiency (Toews et al. 2014).

However, it was not known if, in addition to mtDNA introgression, portions of the nuclear genome also introgressed (Milá et al. 2011, Brelsford et al. 2011, Toews et al. 2014). Given the complex co-evolution that occurs between the nuclear and mitochondrial genomes—particularly between gene products that interact directly in the mitochondria—Toews et al. (2014) predicted that, if nuclear regions had concomitantly introgressed, they would likely contain nuclear mitochondrial gene products. Our WGS data revealed three previously unidentified regions that were moderately differentiated (*F*_*ST*_ > 0.2) between Audubon’s warblers that had mitochondrial haplotypes from the introgressed northern clade versus those from the ancestral southern clade. Of note, the two most divergent regions were in adjacent windows on chromosome 12. Importantly, this region includes the gene *NDUFAF3*, which is a nuclear-encoded mitochondrial gene that encodes an assembly protein in complex I of the electron transport chain (Saada et al. 2009). This differentiation between Audubon’s warblers that differ only in their mitochondrial type suggests that these regions may have co-evolved with mitochondrial gene products and introgressed in tandem with the mitochondrial genome. The PCA (Figure 5D) also provides indirect evidence of introgression: with all three groups included (*S. c. coronata* and both mitochondrial types of *S. c. auduboni*), along PC2—the axis that most strongly separates *S. c. auduboni* with N versus S mitochondrial clades—the northern mtDNA *S. c. auduboni* individuals cluster with *S. c. coronata*, consistent with gene flow in this region for this pair.

We note that, in this case, *F*_ST_ for this region of interest is not high (*F*_ST_ = 0.28). However, to obtain the most power to detect meaningful nuclear differences between the mitochondrial types—as opposed to neutral differences generated by geographic population structure—we sampled warblers from the same locality at the center of the mitochondrial contact zone (Figure 1 orange and red circles). This means that any differences are likely to be smaller than if samples had been drawn from more geographically distant populations on either side of the mtDNA cline. Moreover, the mtDNA cline in this region of the Southwestern USA, while narrower than expected under purely neutral diffusion, is not nearly as narrow as the hybrid zone between *S. c. coronata* and *S. c. auduboni* in the Rockies (Brelsford and Irwin 2009). Thus, selection against mismatched nuclear-mitochondrial types is predicted to be low or moderate, reducing the power to detect associations. Thus, further study and additional sampling along the mitochondrial cline is needed to more definitively conclude whether this region on chromosome 12 introgressed along with mtDNA.

We know of only one other system—Iberian hares—that used WGS to examine an instance of mitochondrial introgression, where mtDNA introgressed from *Lepus timidus* into *L. granatensis* (Seixas et al. 2018). By explicitly testing for adaptive introgression, Seixas et al. (2018) showed that introgression was widespread across the genome, with overrepresentation of introgressed genes involved in spermatogenesis, as well as six mito-nuclear genes that showed high frequency introgression. However, their results more generally suggest a demographic pattern of invasive replacement, with male-biased migration contributing to the strong mito-nuclear discordance, in contrast to explicit adaptive introgression of mtDNA and associated mito-nuclear genes.

## Conclusion

We have been able to contrast the landscapes of divergence using three different methods within the same species complex, a species where previous studies have found a heterogeneous pattern of differentiation. We show that WGS recovered the highest resolution pattern of differentiation, at an equal or better value compared to reduced-representation approaches. We also demonstrate the benefit of this higher resolution for phylogeographic questions by identifying for the first time a small region of the nuclear genome that may have introgressed along with mitochondrial DNA in the Audubon’s warbler. We suggest that further sampling will be needed to confirm this putative instance of nuclear -mitochondrial introgression. However, it showcases the clear benefits and value of WGS approaches compared to reduced representation methods for molecular ecological applications.

## ACKNOWLEDGEMENTS

The authors would like to thank the Museum of Southwestern Biology and the Cornell Museum of Vertebrates (Charles Dardia, curator) for tissue loans. We thank C. Gregory Schmitt, C. Jonathan Schmitt, Cole J. Wolf, Nicholas K. Fletcher, and Borja Milá for facilitating the sample collection. The authors also thank Darren Irwin and the Irwin Lab at the University of British Columbia, where the original GBS data were generated; and Irby Lovette and the Lovette lab at the Cornell Lab of Ornithology, where the original ddRAD and WGS data were generated. The authors would also like to thank Andrew Foote, Evelyn Jensen, Rebecca Taylor, and David Coltman for the invitation to contribute to this special issue of *Molecular Ecology*, as well as three anonymous reviewers whose comments greatly improved the manuscript. Funding was supported by Pennsylvania State University and startup funds from the Eberly College of Science and the Huck Institutes of the Life Sciences.

## DATA ACCESSIBILITY

All data from the present study is published and publicly available. The ddRAD dataset is available at DataDryad, doi: 10.5061/dryad.hs07534. The GBS dataset is available at the NCBI SRA under BioProject # PRJNA471352. The WGS data is available at the NCBI SRA under BioProject# PRJNA630247. The warbler reference genome is available at NCBI BioProject # PRJNA325157.

## AUTHOR CONTRIBUTIONS

Designed research: SS, DPLT, CW, and AB

Performed research: SS, DPLT

Contributed new reagents or analytical tools: DPLT

Analyzed data: SS and DPLT

Wrote the paper: SS, DPLT, CW, and AB

## Notes

### Competing Interest Statement

The authors have declared no competing interest.

