## Supplemental Figures for "Comparing divergence landscapes from reduced-representation and whole-genome re-sequencing in the yellow-rumped warbler (*Setophaga coronata*) species complex"

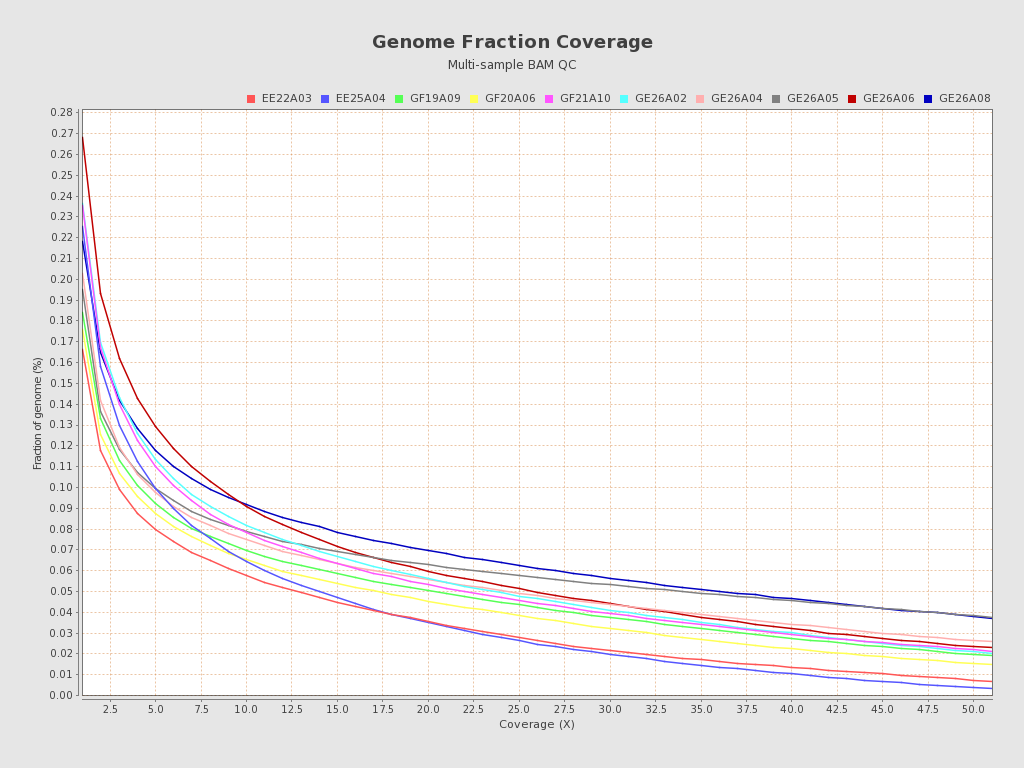

**Figure S1. Distribution of coverage across the genome for ddRAD-seq dataset.** Each line represents one sample.

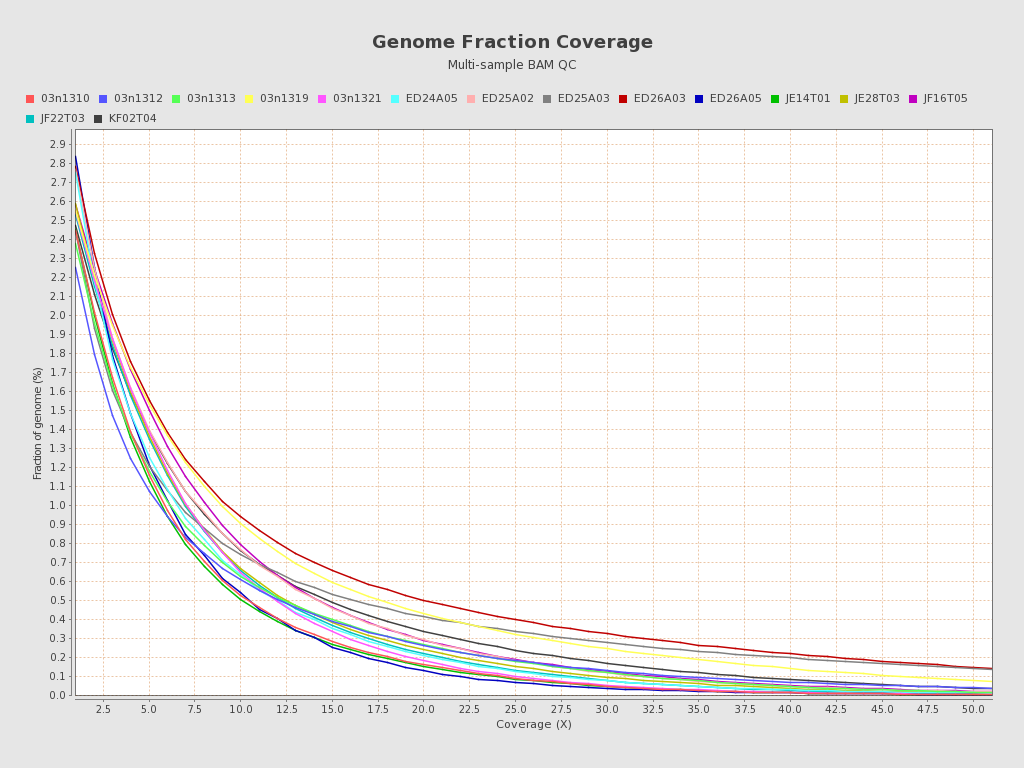

**Figure S2. Distribution of coverage across the genome for GBS dataset.** Each line represents one sample.

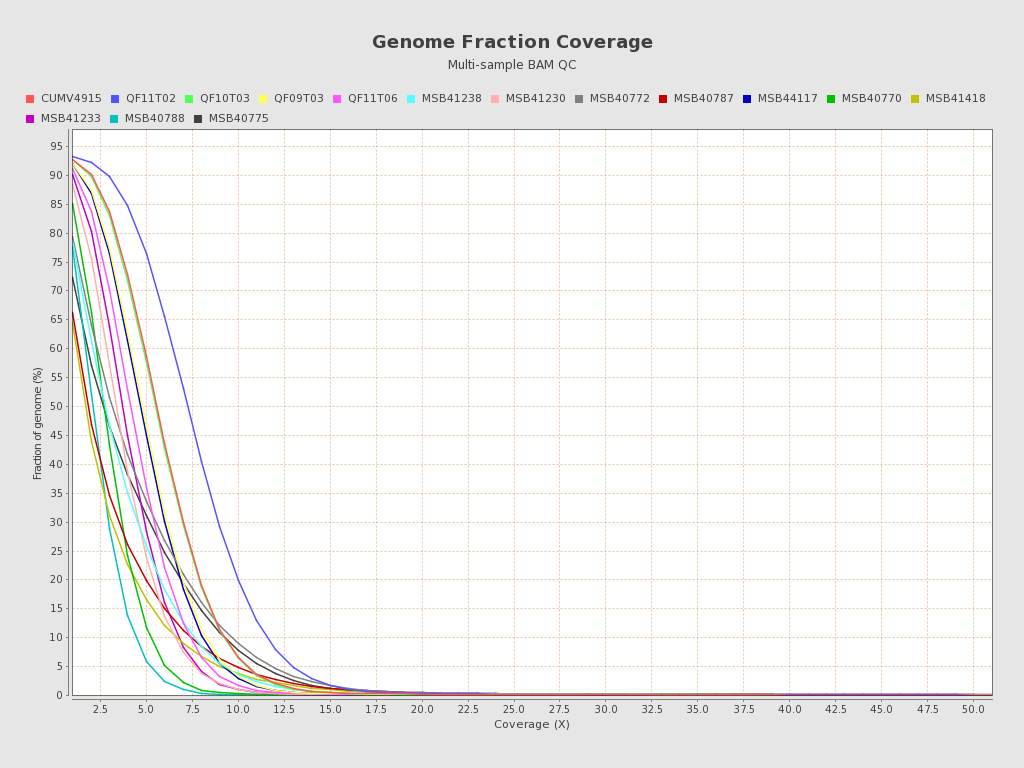

**Figure S3. Distribution of coverage across the genome for WGS dataset.** Each line represents one sample.
